# Complex hybridization between deeply diverged fish species in a disturbed ecosystem

**DOI:** 10.1101/2022.10.08.511445

**Authors:** Shreya M. Banerjee, Daniel L. Powell, Benjamin M. Moran, Wilson F. Ramírez-Duarte, Quinn K. Langdon, Theresa R. Gunn, Gaby Vazquez, Chelsea Rochman, Molly Schumer

**Affiliations:** Department of Biology, Stanford University; Centro de Investigaciones Científicas de las Huastecas “Aguazarca”, A.C.; Center for Population Biology, University of California, Davis; Department of Ecology and Evolutionary Biology, University of Toronto; Hanna H. Gray Fellow, Howard Hughes Medical Institutes

**Author notes:** co-first authorship (Shreya M. Banerjee, Daniel L. Powell).

## Abstract

Over the past two decades researchers have documented the extent of natural hybridization between closely related species using genomic tools. Many species across the tree of life show evidence of past hybridization with their evolutionary relatives. In some cases, this hybridization is complex – involving gene flow between more than two species. While hybridization is common over evolutionary timescales, some researchers have proposed that it may be even more common in contemporary populations where anthropogenic disturbance has modified myriad aspects of the environments in which organisms live and reproduce. Here, we develop a flexible tool for local ancestry inference in hybrids derived from three source populations and describe a complex, recent hybridization event between distantly related swordtail fish lineages (*Xiphophorus*) and its potential links to anthropogenic disturbance.

**Impact Summary:** As sequencing tools have advanced, we have found that barriers between animal species are more porous than once thought. Researchers have found evidence for hybridization between species throughout many branches of the tree of life. In some cases, these hybridization events can involve more than two species. Here, we develop a flexible and user-friendly tool that can be used to identify three-way hybrids and report the discovery of hybrids with ancestry from three swordtail (*Xiphophorus*) species from an anthropogenically impacted site on the Río Calnali in Hidalgo, Mexico. Researchers have studied hybrids between two *Xiphophorus* species along this river for decades, but this is the first documented case of hybridization involving three species. We explore hypotheses for what drove this hybridization event, including anthropogenic pollutants and reduced water quality.

## Introduction

Hybridization, or genetic exchange between species, is common in diverse organisms across the tree of life, and can have important evolutionary consequences (Moran *et al*., 2021). The genetic, ecological, and evolutionary outcomes of hybridization are varied, from facilitating rapid adaptation to exposing genetic incompatibilities. Examples from the recent literature include introgression as a source of genetic rescue (Oziolor *et al*., 2019) and hybridization resulting in decreased tolerance of thermal stressors (Payne *et al*., 2022). While evidence of ancient introgression in the genomes of diverse taxa suggests that hybridization is common in the evolutionary history of many species (Moran *et al*., 2021), a growing number of studies point towards anthropogenic disturbance as contributing to the formation of new hybrid zones (Fisher *et al*., 2006; Kelly *et al*., 2010; Pampoulie *et al*., 2020). As humans modify habitats, there are numerous mechanisms by which anthropogenic environmental disturbance can cause hybridization. These include phenological changes (Chunco, 2014; Vallejo-Marín & Hiscock, 2016), species introductions (Oziolor *et al*., 2019), habitat alterations that lead to new contact zones (Kelly *et al*. 2010), and decreased encounter rates of conspecifics (Willis *et al*., 2011). Environmental disturbance, e.g., pollution with urban effluents and/or reduced water quality, can also directly impact sensory communication, disrupting signals used in mate choice (Seehausen *et al*., 1997; Fisher *et al*., 2006; Powell *et al*., 2022).

Hybridization between pairs of species has been intensively studied for several decades, but a growing body of literature highlights that hybridization events can be complex, involving three or more species (Heliconius Genome Consortium 2012; Toews *et al*. 2018; Langdon *et al*. 2019; Grant and Grant 2020, Natola *et al*. 2022). These types of complex hybridization events are likely to be more common in groups where many species are interfertile and have overlapping ranges. The evolutionary consequences of these events are not as well understood. One possible outcome is “conduit” introgression, where genetic exchange can occur between species that are not in geographic contact through hybridization with a third species (Langdon *et al*. 2019; Grant and Grant 2020; Natola *et al*. 2022). Such dynamics could explain observations in the empirical literature such as cases where gene flow is inferred between geographically isolated species (Cui *et al*., 2013); although there are other potential causes of these patterns including introgression from a now extinct lineage (Ottenburghs, 2020).

*Xiphophorus* species and their hybrids from the Sierra Madre Oriental of eastern Mexico have been intensively studied over two decades. Much of this work has focused on hybridization between two sister-species from the “Northern swordtail” clade, *X. birchmanni* and *X. malinche* (Fig. 1). Throughout its range *X. birchmanni* is sympatric with a distantly related species in the “platyfish” clade, *X. variatus* (Fig. 1). *X. variatus* is also common at many sites where *X. birchmanni* x *X. malinche* hybrids (i.e. Northern swordtail hybrids) are found, but does not reach the high elevations inhabited by pure *X. malinche*. Genetic and historical estimates indicate that hybridization has been occurring between *X. birchmanni* and *X. malinche* in the Río Calnali for more than 40 generations (Rosenthal *et al*., 2003; Schumer *et al*., 2014, 2017). However, despite extensive collections in regions where they co-occur over the last two decades, no hybrids between *X. variatus* and *X. birchmanni* or *X. variatus* and *X. birchmanni* x *X. malinche* have been reported.

**Figure 1.**
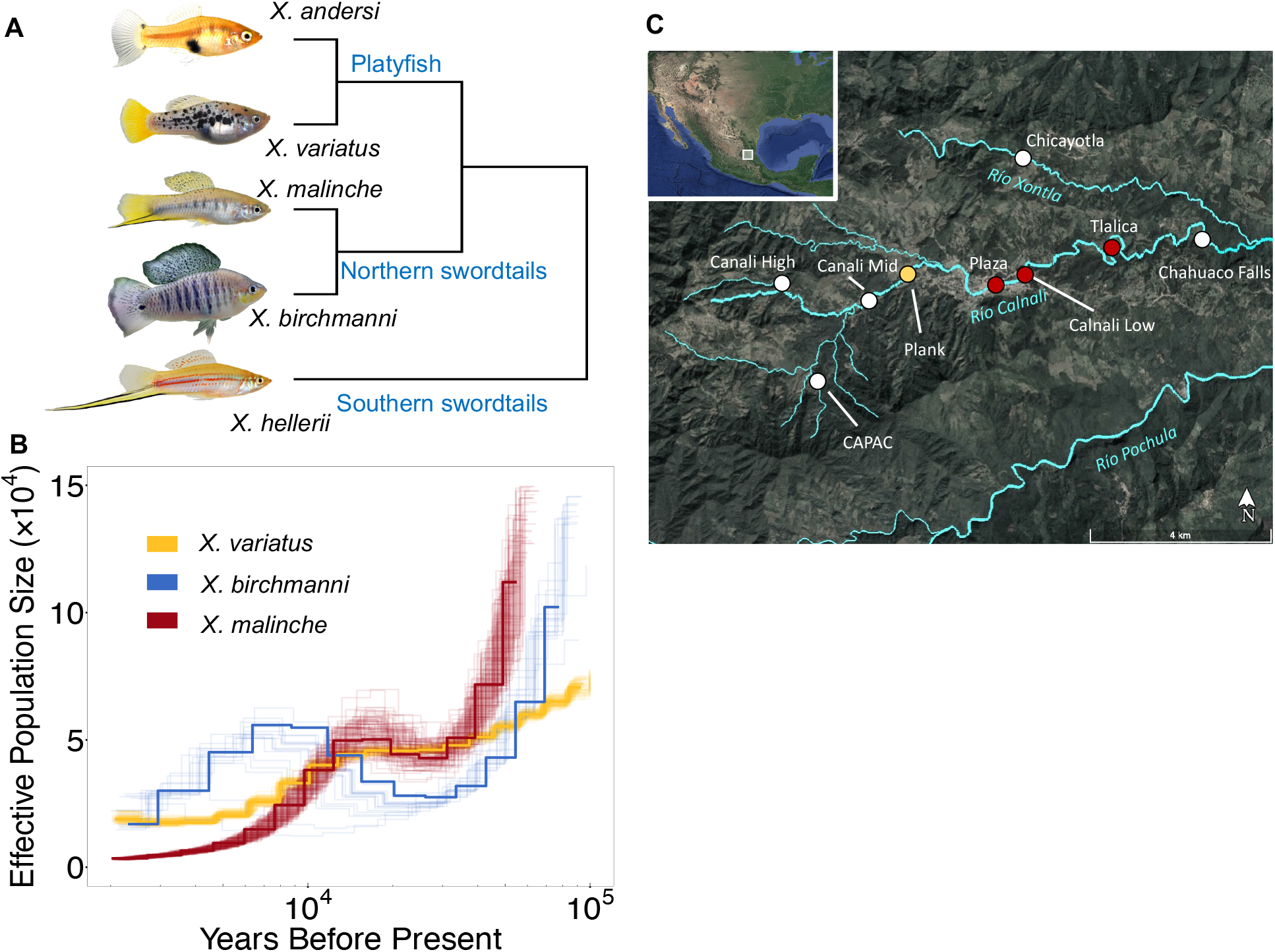
**A**) Simplified phylogeny of the genus *Xiphophorus. X. birchmanni* and *X. malinche* are sister species in the Northern swordtail clade and *X. variatus* belongs to the distantly related platyfish clade. **B**) PSMC results analyzing population history from a single whole-genome sample of *X. variatus* and previously collected data from *X. birchmanni* and *X. malinche* (from Schumer *et al*. 2018). Analysis was conducted with a π/θ ratio of 2, generation time of two generations per year, and mutation rate of 3.5 × 10^−9^. Faint lines reflect bootstrap resampling of the data. **C**) Map of collection sites along the Río Calnali with the focal sites where three-way hybrids have been collected highlighted in red, and upstream site used for comparison in water quality and chemistry samples highlighted in yellow. Inset shows location of *X. birchmanni* x *X. malinche* hybrid populations in Hidalgo, Mexico relative to a map of North and Central America. Images adapted from Google Earth.

Here, we characterize a newly discovered three-way hybridization event involving *X. birchmanni* x *X. malinche* hybrids and *X. variatus* at an anthropogenically disturbed site on the Río Calnali (hereafter the Tlalica site; Fig. 1). To facilitate this analysis, we develop an easy to use and accurate extension of the *ancestryinfer* pipeline (Schumer *et al*., 2020) that enables local ancestry inference of individual hybrids formed from three source populations. We initially identified three-way hybrids based on morphology, and confirmed our observations with whole genome-sequencing and local ancestry inference. We also characterized water quality and chemistry (relevant to the visual and olfactory environment) at Tlalica and other sites along the Río Calnali to explore relationships between environmental disturbance and hybridization. Our results hint at a connection between anthropogenic disturbance and hybridization in these deeply diverged species with a long history of reproductive isolation in sympatry.

## Methods

### Morphological evidence of hybridization between X. variatus, X. birchmanni, and X. malinche

Three-way hybrids were first identified at the Tlalica site based on their unusual phenotype combinations. Male *X. variatus, X. birchmanni*, and *X. malinche* differ in several traits (females are phenotypically similar in many *Xiphophorus* species). *X. malinche* has a modification of the caudal fin known as the “sword” which is absent in the two other species and *X. birchmanni* has a much larger dorsal fin compared to the other two species (Fig. 1). *X. variatus* is characterized by a distinctive diamond body shape (Fig. 1A, 2A), and two horizontal stripes composed of melanophores bracketing the lateral line. By comparison, *X. malinche, X. birchmanni*, and their hybrids are more elongated (Fig. 2A) and have a single, broader horizontal stripe. *X. variatus* have polymorphic melanophore tailspot patterns (described in Borowsky, 1980; Culumber & Rosenthal, 2013) that are distinct from the polymorphic melanophore patterns present in some *X. birchmanni* and *X. birchmanni* x *X. malinche* hybrid individuals (Rauchenberger *et al*., 1990; Culumber, 2014; Powell *et al*., 2020).

**Figure 2.**
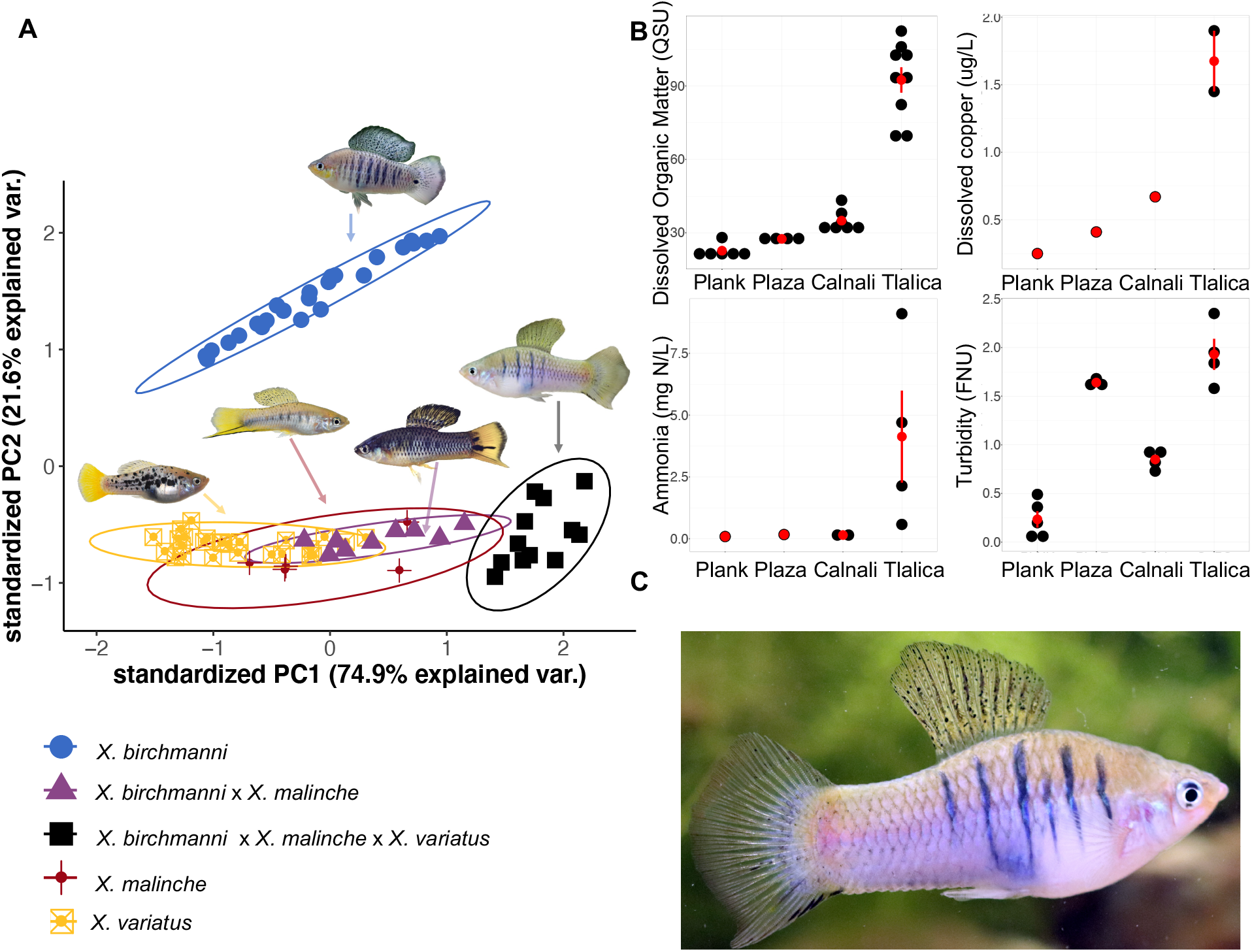
**A**) Principal component analysis of morphology of male *X. malinche, X. birchmanni, X. variatus, X. birchmanni* x *X. malinche* hybrids and confirmed three-way hybrids. **B**) Dissolved organic matter (quinine sulfite units - QSU), ammonia (mg N/L), dissolved copper (ug/L), and turbidity (formazin nephelometric units - FNU) levels measured at Plank, Plaza, Calnali Low (Calnali), and Tlalica in May and June of 2022. Black dots represent independent measurements, red dots represent means, and red bars show one standard error of repeated measurements. **C**) Example of a first generation hybrid individual with 50% *X. variatus* ancestry, 23.8% *X. birchmanni* ancestry, and 26.2% *X. malinche* ancestry. This individual has a short sword, a trait which is unique to *X. malinche*, and large dorsal fin characteristic of *X. birchmanni*, and an overall body shape and vertical barring characteristics of *X. variatus*.

We noticed that two adult males sampled from Tlalica in June 2021 appeared to have *X. variatus*-like characteristics, such as a diamond body shape and dual horizontal stripes, *X. birchmanni*-like characteristics, such as large body size and large rounded dorsal fins, and *X. malinche*-like characteristics, such as short sword extensions (Fig. 2). For these fish and additional male three-way hybrids identified in subsequent collections (n = 12) we measured standard length, body depth, peduncle depth, caudal fin length, dorsal fin width, dorsal fin height, and sword length from photographs of anaesthetized adult male fish using ImageJ (Fig. S1; Schneider, Rasband, & Eliceiri, 2012). We included phenotypes of *X. birchmanni* x *X. malinche* hybrids from a nearby population (Calnali Low; n = 9) and of pure parental species individuals from sites where hybrids have not previously been reported (*X. variatus* from Coacuilco: n = 27, *X. birchmanni* from Coacuilco: n = 24; *X. malinche* from Capac: n = 5) (Fig. 1C). We performed principal component analyses to assess morphological differences between groups.

### Genomic libraries of putative hybrids

For the two male individuals collected in June of 2021 that were morphologically identified as likely three-way hybrids, we produced high-coverage whole genome data (Table S1) following Schumer *et al*. (2016, 2018). Briefly, we extracted DNA from fin clips using half reactions of the Agencourt DNAdvance kit. DNA was sheared by sonication and end-repaired with dNTPs, T4 DNA polymerase, Klenow DNA polymerase, and T4 polynucleotide kinase, then A-tailed with Klenow exonuclease and dATP. Universal Illumina adapters were ligated onto the A-tailed sample using DNA ligase. Samples were purified with the Qiagen PCR Purification kit between steps. Samples were PCR amplified for 12 cycles using the Phusion PCR kit. After amplification, the final PCR product was purified with 18% SPRI beads and sent to Admera Health (South Plainfield, NJ, USA) for sequencing on an Illumina HiSeq 4000.

### Preliminary investigation of three-way hybrids with sppIDer

We used the competitive mapping and read depth analysis pipeline, sppIDer (Langdon *et al*., 2018) as an initial approach to investigate the potential genetic contributions to the two male fish sampled in June of 2021 from Tlalica. We created a combined fasta file by concatenating *X. birchmanni, X. malinche*, and *X. variatus* reference genomes (described in Powell *et al*., 2021). Reads from the high coverage Tlalica males were mapped to this combination reference genome and uniquely mapped reads were used by sppIDer to estimate the proportion of the genome derived from each species (see Langdon *et al*., 2018).

### PSMC and analysis of whole genome sequences

Raw Illumina reads from an *X. variatus* individual from the Coacuilco population (previously sequenced by Powell *et al*., 2020) were aligned to a *de novo* assembly derived from *X. variatus* that was scaffolded with cactus (Armstrong *et al*., 2020) to a chromosome-level *X. birchmanni* assembly (Powell *et al*., 2020). Alignment of reads was performed using *bwa* (Li & Durban, 2009). We used PicardTools and *GATK* (Van der Auwera & O’Connor, 2020) to realign mapped reads around indels and call variant sites in a gvcf format. Individual jobs were run for each chromosome for indel realignment and variant calling. We combined gvcf files for all chromosomes using *bcftools* (Danecek *et al*., 2021). We filtered variant and invariant sites from this combined gvcf file as previously described (Schumer *et al*., 2018). Briefly, we used hard-call thresholds for variant quality scores recommended by GATK and previously validated in swordtails using pedigree data (Schumer *et al*., 2018). For both variant and invariant sites, we masked sites within 5 bp of an INDEL or >2X or <0.5X the average genome-wide coverage. To generate a pseudo-fasta file reflecting variant and masked sites, we used a custom script to generate an insnp file (https://github.com/Schumerlab/Lab_shared_scripts). We used *seqtk* (https://github.com/lh3/seqtk) to generate a new fasta file with variant sites and masked sites updated to reflect the *X. variatus* individual being analyzed.

We next used this data to infer changes in historically effective population size through time using the Pairwise Sequential Markovian Coalescence approach (PSMC, Li & Durbin, 2011). We used a custom script to convert the fasta file to a fastq file (the required input format for PSMC) with uniform quality scores, (https://github.com/Schumerlab/Lab_shared_scripts) and used *seqtk* to exclude scaffolds that did not belong to the 24 major *Xiphophorus* chromosomes. We assumed a mutation rate of 3.5 × 10^−9^ per basepair per generation, a generation time of half a year, and set the -r parameter to 2 (Schumer *et al*., 2018). We compared these results for *X. variatus* to those previously published for *X. birchmanni* and *X. malinche* (Schumer *et al*., 2018). For comparison to our results for *X. variatus*, we included only one sample per species.

### Low-coverage whole genome sequencing of individuals collected from Tlalica and nearby sites

We extracted DNA from fin clips using the Agencourt DNAdvance bead-based kit as specified by the manufacturer except that we used half-reactions. We prepared tagmentation-based libraries for low-coverage whole genome sequencing as previously described (Payne *et al*. 2022). Briefly, DNA was enzymatically sheared using the Illumina Tagment DNA TDE1 Enzyme and Buffer Kit, amplified in a dual-indexed PCR reaction for 12 cycles, pooled, and bead purified with 18% SPRI magnetic beads. Libraries were sent to Admera Health (South Plainfield, NJ, USA) to be sequenced on a HiSeq 4000.

### Design and performance tests of three-way local ancestry inference

To perform three-way local ancestry inference, we adapted our previously developed pipeline, *ancestryinfer* (which only allowed for two source populations; Schumer *et al*., 2020), to accommodate three reference genomes and source populations. Briefly, we modified the program to detect whether two or three reference genomes were provided in the configuration file (see Appendix 1 – user manual). When three reference genomes are provided, *ancestryinfer* maps reads to all three genomes and identifies and excludes any reads that do not map uniquely to any of the three references. Using the coordinate space of reference genome 1, it tabulates counts for each allele at each ancestry-informative site and runs AncestryHMM in the three source population mode (Corbett-Detig & Nielsen, 2017). Users can optionally provide priors for the number of generations since initial admixture for each source population and priors for admixture proportions from each source population.

We searched for candidate ancestry-informative sites from high coverage whole genome sequences (*X. variatus* n=2, *X. malinche* n=4, *X. birchmanni* n=26; samples from Schumer *et al*., 2018; Powell *et al*., 2020, 2021). Although we use a small number of high coverage samples in this initial step based on available data, we filter sites using a large number of individuals of each species (see below). We identified biallelic sites that differentiated any of the three focal species. Initial analysis suggested that issues with accuracy arise from imbalance in the number of ancestry-informative sites between pairs of species. We thinned to an approximately equivalent number of informative sites between all pairs of species. To do so, we retained all ancestry-informative sites that distinguished *X. birchmanni* and *X. malinche*, and every other site that distinguished *X. variatus* from either of these two species.

We refined this candidate set of ancestry-informative sites using low-coverage population data from each species (*X. malinche* n=28, *X. birchmanni* n=107 – Schumer *et al*., 2018; *X. variatus* n=145 – this study). Note that per basepair heterozygosity is much lower in *X. malinche*, approximately 1/4 of the levels observed in *X. birchmanni* or *X. variatus* (0.0003 per basepair versus ∼0.001 respectively). Low nucleotide diversity in *X. malinche* is attributable to low historical effective population sizes in this species (Schumer *et al*., 2018). Average coverage per individual was ∼1X. Because this is low coverage data, we did not perform explicit variant calling but instead used bcftools mpileup to determine the observed counts for each allele at each candidate ancestry-informative site in the three source populations. We then excluded ancestry-informative sites that did not have equal to or greater than a 90% frequency difference between at least one pair of species (e.g. *X. birchmanni* vs *X. malinche, X. birchmanni* vs *X. variatus, X. malinche* vs *X. variatus*). This resulted in a final set of 997,366 ancestry-informative markers throughout the 750 Mb genome.

Using this set of ancestry-informative sites and estimated parental allele frequencies determined from the individuals described above, we ran *ancestryinfer* on a set of parental individuals that were not used in the training datasets (n_*variatus*_ = 30; n_*birchmanni*_ = 12; n_*malinche*_ = 10) as a first performance check on empirical data. We found that *ancestryinfer* correctly inferred that these individuals were unadmixed and derived from the correct parental population (Fig. S2). We also performed a similar analysis on hybrids from *X. birchmanni* x *X. malinche* hybrid populations that are allopatric with respect to *X. variatus* (Totonicapa, n=30, and Tlatemaco, n= 23; Fig. S3). See Supporting information 1 for additional performance testing and simulations.

We note that we do not have access to any populations in which *X. birchmanni* does not co-occur with *X. variatus*. Thus, if there was admixture between *X. birchmanni* and *X. variatus* in the *X. birchmanni* source populations that we have failed to detect, our approach could underestimate the degree of contemporary gene flow between these species.

### Local ancestry inference and data processing of three-way hybrids

We proceeded with local ancestry analysis of individuals collected from the Tlalica population (n=64) and previously collected samples from upstream (n=553; Table S2) and downstream (n=25) of this site on the Río Calnali. We also ran *ancestryinfer* on all sequenced pure *X. birchmanni* and *X. variatus* from Coacuilco, an allopatric site in a different drainage, to confirm their ancestry (n=745; samples from Powell *et al*., 2020 and this study; Table S3). For hybrid individuals from the Río Calnali, we provided priors for admixture proportions from the three source populations based on sppIDer results. *ancestryinfer* accepts priors for the time since initial admixture for all source populations (see Corbett-Detig & Nielsen, 2017). Based on past results for *X. birchmanni* x *X. malinche* (Schumer *et al*. 2014, 2017) and the results of an initial run of *ancestryinfer* without specifying a prior for admixture time, we set the prior admixture time between *X. malinche* and *X. birchmanni* to 50 and the prior admixture time between this admixed population and *X. variatus* to 2. We excluded individuals with fewer than 500,000 reads, based on previous simulation results that indicated accuracy of local ancestry inference is reduced in individuals with <0.2X coverage (Schumer *et al*., 2020). This analysis resulted in posterior probabilities for each of the six possible ancestry states (homozygous *X. birchmanni*, homozygous *X. malinche*, homozygous *X. variatus* and each possible heterozygous combination) at 900,343 ancestry-informative sites throughout the genome.

We used a posterior probability threshold of 0.9 to convert ancestry probabilities to hard-calls. For ancestry-informative sites that did not have a probability of ≥0.9 for any ancestry state, we converted the probabilities for those sites to NA. The average level of missing data in three-way hybrid individuals after imposing this hard-call threshold was 0.03%.

### Water quality and chemistry at Tlalica

Tlalica is ∼1 km away from the municipal landfill of the city of Calnali and 2.7 km downstream from the outfall of a sewage treatment plant. During the wet season, we observed a small tributary running through the municipal landfill into the Río Calnali approximately 250 meters upstream of Tlalica. On one sampling occasion, we observed sewage effluent flowing into the river from the treatment plant upstream of Tlalica. On another occasion a break in the sewer line upstream of the treatment plant led to contamination of the river ∼3 km upstream of the sampling site. Accordingly, we expected water quality at the Tlalica site to be lower than upstream sites (Fig. 1), and hypothesize that this could contribute to the hybridization observed between distantly related *Xiphophorus* species.

We collected water samples in May and June of 2022 at a relatively undisturbed upstream site (Plank), and at the three sites where we found genetic evidence of one or more three-way hybrids (see Results; Plaza, Calnali Low, and Tlalica; Fig. 1). All focal sites contained both *X. variatus* and *X. birchmanni* x *X. malinche* hybrids. We measured fluorescent dissolved organic matter (DOM) and turbidity using an EXO2 multiparameter sonde (YSI, Yellow Springs, OH).

We used a 9300 colorimeter (YSI, Yellow Springs, OH) to quantify ammonia. We quantified concentrations of dissolved copper (using a 0.45 μm polyethersulfone membrane, and acidification to pH ∼2.0 with trace metal grade nitric acid) in water using inductively coupled plasma mass spectrometry (ICP-MS) by following the modified version of the APHA3030B/6020A methods. See Supporting information 2 for additional water quality and chemistry metrics collected.

## Results

### Morphological results and demographic survey of the Tlalica population

Three-way hybrids were morphologically distinct from pure parental individuals and from *X. birchmanni* x *X. malinche* hybrids found in nearby populations (Fig. 2, Table S4). They were most morphologically distinct from other groups analyzed along PC1 (74.9% of variation explained), and clustered with *X. malinche, X. variatus*, and Northern swordtail hybrid individuals along PC2 (21.6% of variation explained; Fig. 2).

Based on visual phenotypes, a large majority of individuals collected from the Tlalica site in May 2021, November 2021, February 2022, and May 2022 were classified as *X. variatus*. From visual phenotypic data alone, *X. variatus*-like individuals outnumbered *X. birchmanni* x *X. malinche* hybrids by ∼33:1 (based on 571 individuals collected in May 2022). Genotype data from a subset of fish collected at Tlalica that were categorized as *X. variatus* indicate that we can accurately differentiate them based on morphology alone (Fig. S4).

### *History of divergence between X. variatus, X. birchmanni*, and *X. malinche*

*X. variatus* and the Northern swordtail clade to which *X. birchmanni* and *X. malinche* belong are deeply divergent (Fig. 1; Schumer *et al*. 2014; 2016; 2018). Pairwise sequence divergence between *X. birchmanni – X. variatus* and *X. malinche – X. variatus* is 1.42% and 1.43% respectively. Because *X. malinche* has undergone a severe recent bottleneck (see below, Fig. 1), we focus on comparisons between *X. birchmanni* and *X. variatus* here. The per-site heterozygosity (θ_π_) for *X. variatus* is 0.11%, similar to that observed in *X. birchmanni* (0.12%; Schumer *et al*., 2018). Assuming that the ancestral θ is close to that of *X. birchmanni* and *X. variatus*, we estimate the divergence time between the two clades is approximately 7.5 in units of 4*Ne* generations (using the relationship T_div/2*Ne*_ = D_xy_/θ - 1).

Comparing PSMC results for *X. variatus* to those previously inferred for *X. birchmanni* and *X. malinche* highlights differences in the inferred effective population sizes of each species over time (Fig. 1). We estimated the long-term effective population size of *X. variatus* from one individual to be approximately 50,000 individuals, similar to our previous estimates for *X. birchmanni* (48,000–53,000; Powell *et al*., 2021). However, the timing and extent of demographic fluctuations varies between the two species (Fig. 1). *X. malinche* differs more substantially in its inferred demographic history from both *X. birchmanni* and *X. variatus* given the strong bottleneck that has persisted through much of its recent history (Fig. 1; Schumer *et al*., 2018).

Assuming a long-term effective population size of 50,000 individuals, and the divergence time in 4*Ne* generations calculated above, we estimate the divergence time between *X. variatus* and *X. birchmanni* (and *X. malinche*) to be approximately 1.5 million generations.

### Ancestry analysis of Tlalica hybrids and nearby populations

Initial analysis of genomic data with sppIDer indicated that males visually categorized as three-way hybrids were likely hybrids between *X. variatus, X. birchmanni*, and *X. malinche* (Fig. S5). We found that for each Tlalica male 52% of the reads preferentially mapped to the *X. variatus* reference genome, 30-34% mapped to the *X. birchmanni* reference genome, 26-29% mapped to the *X. malinche* reference genome.

This finding led us to develop local ancestry inference for three-way admixture for these species (see Methods). The results of our local ancestry inference analysis indicated that seventeen individuals sequenced from the Tlalica population were early generation hybrids between *X. variatus, X. birchmanni*, and *X. malinche*. Among individuals at Tlalica with Northern swordtail ancestry, we estimate the frequency of three-way hybrids to be ∼10% of individuals; this estimate is based on individuals collected before May 2022 since in later samples we selectively collected suspected three-way hybrids (Fig. 3B; Table S5). Three-way hybrid individuals derived ∼50-75% of their genomes from *X. variatus*.

**Figure 3.**
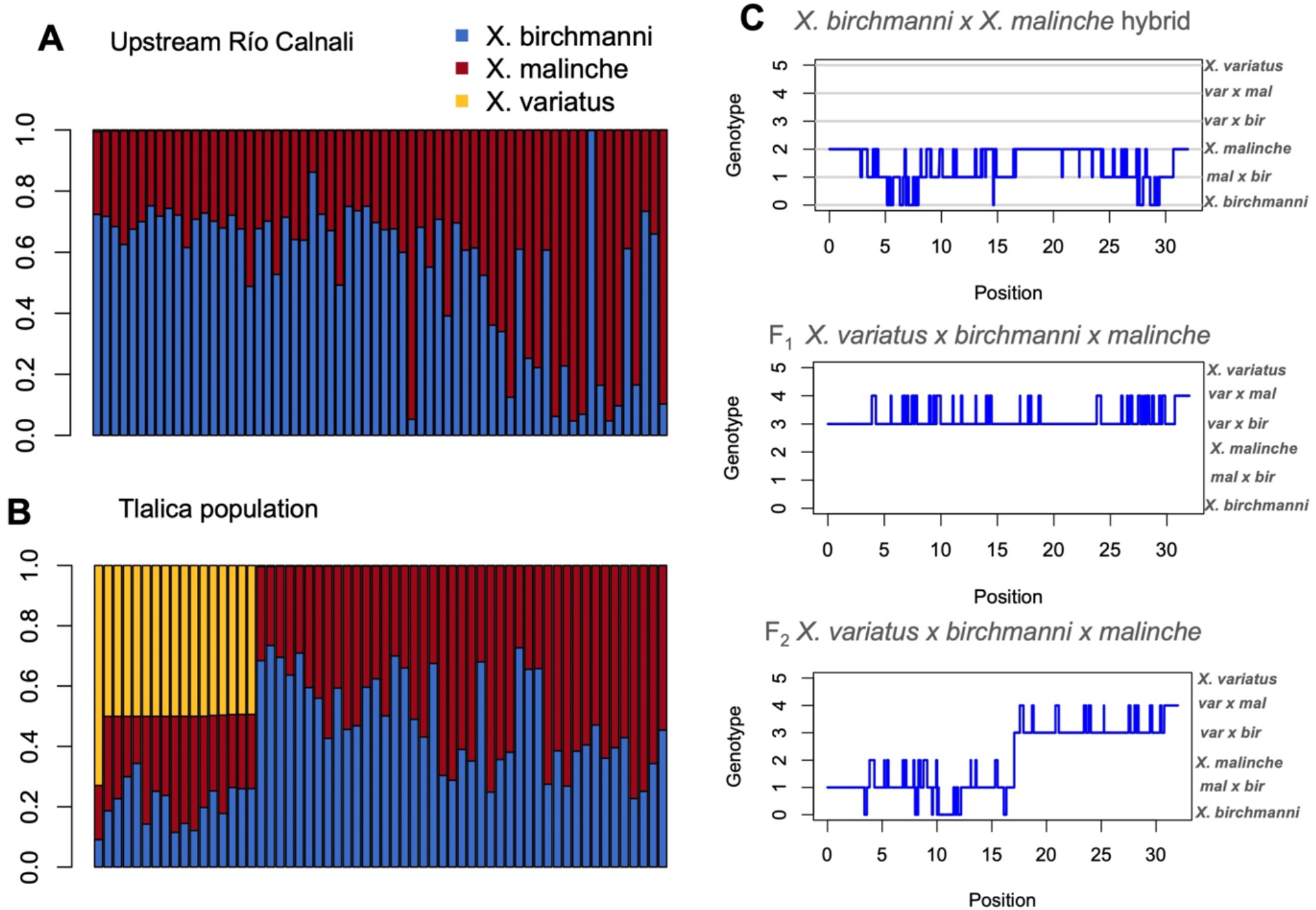
**A**) Distribution of genome-wide ancestry for individuals collected upstream of the Tlalica site on the Río Calnali (N=553, downsampled for visualization to 64). Stacked plots show the estimated proportion of each individual’s genome derived from *X. variatus* (yellow), *X. birchmanni* (blue), and *X. malinche* (red) based on local ancestry inference with *ancestryinfer*. Individuals sampled from upstream sites have near-zero introgression from *X. variatus*. **B**) Distribution of genome-wide ancestry for individuals with suspected swordtail ancestry collected at the Tlalica site on the Río Calnali (N=64). Stacked plots show the estimated proportion of each individual’s genome derived from *X. variatus* (yellow), *X. birchmanni* (blue), and *X. malinche* (red). While some *X. birchmanni* x *X. malinche* hybrid individuals sampled from Tlalica lack *X. variatus* ancestry, a substantial proportion derive some of their genome from *X. variatus*. **C**) Local ancestry inferred along chromosome 1 for individuals of different hybrid types. The genotype on the y-axis corresponds to the ancestry class at that marker: 0 – homozygous *X. birchmanni*, 1 – heterozygous *X. birchmanni* x *X. malinche*, 2 – homozygous *X. malinche*, 3 – heterozygous *X. birchmanni* x *X. variatus*, 4 – heterozygous *X. malinche* x *X. variatus*, 5 – homozygous *X. birchmanni*. In the top plot, a typical *X. birchmanni* x *X. malinche* hybrid from the Río Calnali is shown. In the middle plot, a first generation hybrid between an *X. birchmanni* x *X. malinche* mother and *X. variatus* father is shown (parental source populations inferred based on mitochondrial ancestry, Table S5). Note that across the entire chromosome this individual is either heterozygous *X. birchmanni* x *X. variatus* or heterozygous *X. malinche* x *X. variatus* (genotype classes 3 or 4). In the bottom plot, ancestry for a “backcrossed” three-way hybrid individual is shown. This individual is inferred to be the offspring of a first generation three-way hybrid and a *X. birchmanni* x *X. malinche* mother. Note that this individual is heterozygous for *X. variatus* ancestry over only approximately half of its chromosome.

Samples collected from Tlalica that did not show evidence of ancestry derived from all three species fell into two categories: hybrids between *X. birchmanni* and *X. malinche* and pure *X. variatus* (Fig. 3). Hybrids between *X. birchmanni* and *X. malinche* derived approximately 25-75% of their genomes from either of these parent species (Fig. 3). This is consistent with admixture proportions observed in hybrids between *X. birchmanni* and *X. malinche* at sites up and downstream of Tlalica (Fig. 1; Fig. 2; Schumer *et al*., 2017). This suggests that three-way hybrids with *X. variatus* originated from admixture with already extant *X. birchmanni* x *X. malinche* hybrids. Indeed, *X. birchmanni* and *X. malinche* ancestry tract lengths in three-way hybrids are similar to those observed in *X. birchmanni* x *X. malinche* hybrids at nearby sites (average minor parent ancestry tract length ∼150-200 kb; Schumer *et al*., 2017). Samples preliminarily categorized as *X. variatus* based on morphology from the Tlalica population show no evidence of introgression from *X. birchmanni* or *X. malinche* (Fig. S4).

Given the proximity of Tlalica to previously sampled sites on the Río Calnali (∼3 km), and the fact that *X. variatus* is sympatric with several *X. birchmanni* x *X. malinche* hybrid populations along the river (Fig. 1, Fig. 3), we asked if there is evidence of three-way hybridization at other sites. We performed three-way local ancestry inference on 578 historically and newly collected samples from other sites on the Río Calnali from 2003 to 2022 (previously assumed based on morphology to represent *X. birchmanni* x *X. malinche* hybrids; Schumer *et al*., 2017; Table S2). We identified only two three-way hybrids from these sites — a female from the Plaza site who derived ∼25% of her genome from *X. variatus* and a male from Calnali Low who derived 50% of his genome from *X. variatus* (Fig. 3; 0.4% of sequenced specimens from other sites).

### Inference about the generation of admixture using ancestry tract lengths

Observed admixture proportions for three-way hybrids (25-75% *X. variatus* ancestry across all samples) suggests that these individuals might be early generation hybrids between *X. variatus* and Northern swordtail hybrids. Some of these samples are clearly first generation hybrids between *X. variatus* and *X. birchmanni* x *X. malinche* hybrids based on local ancestry patterns (N=17; Fig. 3). These individuals derived 50% of their genomes from *X. variatus* and were heterozygous for *X. variatus* ancestry at nearly every ancestry-informative site across the genome (>99.5% across individuals; Fig. 3). The few sites inferred to be homozygous *X. variatus* are consistent with our expected error rate (see Methods).

Two individuals with substantial *X. variatus* ancestry did not have ancestry patterns consistent with those expected for first generation hybrids. Both their observed admixture proportions (25% and 75% *X. variatus*, respectively) and the lengths of ancestry tracts heterozygous or homozygous for *X. variatus* ancestry indicate that these individuals are likely backcrosses between a three-way hybrid with a pure *X. variatus* individual (Fig. 3). The identification of two second-generation three-way hybrids indicates that hybrids between *X. birchmanni, X. malinche*, and *X. variatus* are at least partially fertile.

Since the mitochondrial genome is maternally inherited, it allows us to infer the likely maternal ancestry for the three-way hybrids identified. All three-way hybrids (n = 19) sequenced had mitochondrial ancestry derived from either *X. birchmanni* or *X. malinche* (Table S5), suggesting that the mothers of all three-way hybrid individuals sampled to date were *X. birchmanni* x *X. malinche* hybrids. Skews in maternal ancestry may be the result of population demography, differences in the strength of mate discrimination across groups, or impacts of cross direction on the viability of hybrids.

### Water quality and chemistry analysis

Dissolved organic matter, ammonia, dissolved copper, and turbidity were all elevated at the Tlalica sampling site in comparison with the relatively undisturbed upstream Plank site when we collected water samples in the spring of 2022 (Fig. 2B). Two sites between Plank and Tlalica where three-way hybrids were detected at low frequencies, Plaza and Calnali Low, had intermediate values of dissolved organic matter, ammonia, and dissolved copper (Fig. 2B). With one season of data collection, we focus on qualitative patterns in the results. However, our results show a pattern of elevated pollution and turbidity at sites with higher frequencies of hybridization between *X. variatus* and *X. birchmanni* x *X. malinche* hybrids. Additional water quality and chemistry metrics are shown in Fig. S6.

## Discussion

Here we characterize wild caught *Xiphophorus* individuals with ancestry from three parental species — *X. variatus, X. birchmanni*, and *X. malinche* — using multiple approaches. Though *X. birchmanni* x *X. malinche* hybrids were first reported nearly two decades ago (Rosenthal *et al*. 2003), hybridization with *X. variatus* has not been previously reported. These results are remarkable given that *X. variatus* have ∼1.5% sequence divergence from the Northern swordtail clade. This is similar to divergence between chimpanzees (*Pan troglodytes*) and gorillas (*Gorilla spp*.) (Chen & Li, 2001), highlighting the unusually deep nature of this hybridization event.

To facilitate this work, we developed a user-friendly pipeline for running local ancestry inference in hybrids which derive their genomes from three source populations, as well as a collection of simulation scripts to test expected performance (see Appendix 1). Although several methods have been developed that accommodate local ancestry inference with three source populations (reviewed in Wu *et al*. 2021), there are few pipelines available that allow researchers to move from raw reads to probabilities of ancestry across the genome. By expanding our previously developed local ancestry inference pipeline and simulation scripts (Schumer *et al*., 2020), we are able to provide a toolkit that can be used by researchers to study complex hybridization events in diverse species groups.

The existence of natural hybrids between *X. variatus* and *X. birchmanni* x *X. malinche* hybrids is surprising. Despite extensive collections over the past two decades and substantial range overlap between *X. variatus, X. birchmanni*, and *X. birchmanni* x *X. malinche* hybrids, there are no reports of contemporary hybridization involving *X. variatus*. However, past work has identified a small genomic contribution from the lineage leading to *X. variatus* to the ancestors of *X. birchmanni* and *X. malinche* (Schumer *et al*. 2018), which indicates that gene flow has occurred historically. This event contributed ∼2-4% of the genome to present-day *X. birchmanni* and *X. malinche* (Schumer *et al*. 2018). Together with our present data, this suggests that hybridization between these groups is possible, albeit rare.

*X. variatus* are sympatric with natural *X. malinche* x *X. birchmanni* hybrid populations at several upstream sites along the Río Calnali where we have found no evidence of three-way hybridization. In our analyses of 642 Northern swordtail individuals from sites along the river, we have identified fifteen individuals with three-way hybrid ancestry at Tlalica, one individual at Plaza, and one individual at Calnali Low. Moreover, *X. birchmanni* and *X. variatus* are sympatric over much of *X. birchmanni*’s range, but there has been no evidence of hybridization outside of the three-way hybrids reported here (Kallman & Kazianis 2006).

The lack of evidence for contemporary hybridization involving *X. variatus, X. birchmanni* or *X. birchmanni* x *X. malinche* hybrids outside of the Tlalica site and nearby sites suggests that something is unique about this locality. We predict that the demography of the Tlalica community and its disrupted water quality and chemistry play an important role in this unusual hybridization event. Anthropogenic disturbance via wastewater effluent and landfill leachate could have facilitated hybridization via two mechanisms: 1) by decreasing the abundance of *X. birchmanni* x *X. malinche* hybrids with respect to *X. variatus* and 2) by disrupting sensory cues used in mate choice.

*X. birchmanni, X. malinche*, and their hybrids are more sensitive to poor water quality than *X. variatus* (Mercado-Silva *et al*., 2006; personal observation), and *X. variatus* individuals vastly outnumber *X. birchmanni* x *X. malinche* hybrids at the Tlalica site. Notably, nowhere else on the Rio Calnali have we observed such a strong demographic skew toward *X. variatus*. Past research has suggested that female mate preferences can weaken when the density of conspecifics is low and encounter rate with heterospecifics is high, or the search cost of finding a conspecific mate is very high (Cotton *et al*. 2006; Lehmann 2007; Verzijden *et al*. 2011; Stoffer & Uetz 2015; Delclos *et al*. 2020), including in *X. birchmanni, X. malinche*, and *X. variatus* (Fisher & Rosenthal, 2010). This raises the possibility that female *X. birchmanni* x *X. malinche* hybrids mated with *X. variatus* males because there were so few Northern swordtail mates available. This hypothesis is further supported by the lack of *X. variatus* mitochondrial ancestry in three-way hybrids (Table S5). In *Xiphophorus*, females tend to have much stronger conspecific mate preferences than males (Rosenthal & Garcia de Leon, 2011) and *X. variatus* females in the Tlalica population have access to many conspecific males.

In addition to demography, our results are consistent with a role of anthropogenic shifts in water quality and chemistry in this rare hybridization event. In many *Xiphophorus* species, female mate choice is driven in large part by species-specific olfactory signals (Crapon De Caprona and Ryan 1990; McLennan and Ryan, 1999; Wong *et al*. 2005; Fisher & Rosenthal 2006; Rosenthal *et al*. 2011; Verzijden *et al*. 2011). Research has shown that organic and inorganic substances can alter the ability of female *X. birchmanni* to distinguish conspecific from heterospecific males (Fisher, Wong, & Rosenthal, 2006, Powell *et al*. 2022). Levels of several chemicals observed at Tlalica may be sufficient to disrupt olfactory communication and drive hybridization between *X. variatus* and *X. birchmanni* x *X. malinche* hybrids. Elevated dissolved organic matter has been shown to impair chemical and/or visual communication in some fish species at concentrations of ∼1 mg/L of humic acid (Hubbard *et al*., 2002; Mobley *et al*., 2020), similar to concentrations found in Tlalica and Calnali (Table S6). Ammonia can impair generation of electric impulses in neurons and lead to health effects at low concentrations (1.3-3.5 mg N/L; Ip, Chew, and Randall 2001). Because ammonia was found at concentrations above toxic thresholds at Tlalica (Fig. 2), future research should test hypotheses regarding whether ammonia affects mating behavior. Finally, the copper concentrations detected at Tlalica were similar to the concentrations reported to disturb olfactory perception and olfactory-mediated behaviors in other fish species (∼2 μg/L; Sandahl *et al*., 2007; Morris *et al*., 2019).

Although *Xiphophorus* respond most strongly to olfactory sexual signals, visual cues are also important in mating decisions (Crapon de Caprona and Ryan, 1990; McLennan and Ryan, 1999; Fisher *et al*., 2006b; Verzijden and Rosenthal 2011; Delclos *et al*. 2020). Thus, the increased turbidity of water at Tlalica and nearby sites may also play a role in the breakdown of reproductive barriers (Seehausen et al. 1997). Turbid waters could indirectly facilitate hybridization among *X. variatus* and *X. birchmanni* x *X. malinche* by disturbing the transmission of visual cues involved in species/mate recognition. Testing hypotheses about the impacts of water chemistry on mate choice using both chemical treatments and mate choice trials is an exciting avenue for future investigation (e.g. as in Fisher *et al*., 2006).

What are the consequences of the complex hybridization events that researchers are beginning to uncover? One possible outcome when three species hybridize is the potential for gene flow between two species that would otherwise not contact each other. Such “conduit introgression” has been described in several systems (Heliconius Genome Consortium 2012; Toews *et al*. 2018; Langdon *et al*. 2019; Grant and Grant 2020, Natola *et al*. 2022). While the scenario uncovered at Tlalica is more complex since hybridization is occurring between pure *X. variatus* and *X. birchmanni* x *X. malinche* hybrids, the effects on dynamics of gene flow could be similar. Specifically, because *X. malinche* does not overlap with *X. variatus*, admixture with *X. birchmanni* x *X. malinche* hybrids could be a route for contemporary gene flow between *X. variatus* and *X. malinche*.

## Conclusions

We present evidence of a deep hybridization event between the platyfish and Northern swordtail clade, involving genetic material from three source species. Given the abundance of first-generation three-way hybrids over multiple sampling seasons and the rarity of second generation or later hybrids in our sample, we predict that there are significant costs in terms of viability or fertility of this distant cross. While contemporary hybrids between these species have not been previously reported, ancient hybridization between them has been inferred (Schumer *et al*. 2018). Our data suggest that this unusual hybridization event may be linked to anthropogenic disturbance in the local environment. Disentangling the mechanisms through which anthropogenic disturbance contributes to hybridization in this system in an exciting direction for future work.

## Acknowledgements

We thank Heidi Fisher, Vitor Sousa, Gil Rosenthal, Claudia Bank, and members of the Schumer and Rochman labs for helpful discussion and/or feedback on earlier versions of this work. We are grateful to the Mexican federal government for permission to collect samples (Permiso de Pesca de Fomento no. PPF/DGOPA-064/20). We thank Stanford University and the Stanford Research Computing Center for providing computational support for this project. This work was supported by NSF GRFP 2019273798 to B. Moran, NSF PRFB (2010950) to Q. Langdon, and a Human Frontiers in Science Grant to M. Schumer & C. Rochman (RGY0081). The authors declare no conflicts of interest.

## Author contributions

S.M. Banerjee, D.L. Powell, and M. Schumer conceived of this project. S.M. Banerjee, D.L. Powell, B.M. Moran, T. Gunn, and W.F. Ramírez-Duarte collected data, S.M. Banerjee, D.L. Powell, B.M. Moran, Q. Langdon, W.F. Ramírez-Duarte, and M. Schumer analyzed data. M. Schumer adapted *ancestryinfer* for three-way local ancestry inference. M. Schumer and C. Rochman oversaw the project. All authors wrote the manuscript.

## Data availability

All raw sequencing data for this project will be deposited on NCBI’s SRA. All water quality and chemistry data and ancestry calls will be deposited on Dryad. Computational pipelines and analysis scripts are available at https://github.com/Schumerlab.

## Notes

### Competing Interest Statement

The authors have declared no competing interest.

### Summary of Updates

Correction to figure 3B: individuals collected in spring of 2022 added to plot

