## Supporting Information for "Complex hybridization between deeply diverged fish species in a disturbed ecosystem"

*Supporting Information 1. Performance and sensitivity of the three-way HMM*

We used several approaches to test the accuracy of local ancestry inference with three source populations, including analysis of pure parental individuals (Fig. S2), analysis of known hybrids between two out of three of the source populations (Fig. S3), and analysis of simulated data.

An advantage of simulation-based approaches is that the ground truth of the ancestry state at each site along the genome is known. We designed a simulation pipeline that we applied to simulate *X. birchmanni* x *X. malinche* x *X. variatus* hybrids but can be used as a tool to model complex hybridization in any system. These scripts are described in Appendix 1 and released with this version of *ancestryinfer* (https://github.com/Schumerlab/ancestryinfer).

Briefly, users can model arbitrarily complex hybrid demographic history using the program SELAM (Corbett-Detig & Jones, 2016), which will simulate ancestry tracts along a chromosome for individuals. SELAM can perform simulations in the presence or absence of selection but all simulations performed for this study lacked selection. Our simulation script takes the ancestry tract coordinates generated by SELAM and user-provided reference genomes as input. The pipeline generates simulated chromosomes and simulated Illumina reads for each individual, as well as bed files indicating the true genotype at each site across the chromosome. The number of simulated Illumina reads per individual (i.e. desired coverage) is specified by the user. This simulated Illumina data can then be run through the *ancestryinfer* pipeline using the 3-way ancestry calling option. After running the *ancestryinfer* pipeline, users can summarize accuracy within and across individuals using the provided scripts (Appendix 1).

To evaluate expected accuracy in *X. birchmanni* x *X. malinche* x *X. variatus* hybrids, we performed simulations matching the likely demographic history and ancestry of three-way hybrid individuals. Several analyses suggested that a subset of the individuals sampled from Tlalica were first or early generation hybrids (see Results). Using past results on the inferred demographic history of hybridization between *X. birchmanni* and *X. malinche* in the Río Calnali (Schumer *et al.*, 2017) we used SELAM to simulate an admixture event between two populations representing *X. birchmanni* and *X. malinche* at equal admixture proportions 50 generations ago, and simulated admixture with *X. variatus* two generations before the present. Using these simulated ancestry tracts and the revised *ancestryinfer* pipeline (see Methods), we generated simulated genomes and reads for 100 hybrid individuals. We then ran the *ancestryinfer* pipeline in the three-way calling mode using the same ancestry informative markers used in our analysis of the real data.

Following simulations, we converted posterior probabilities for the six possible ancestry states at each ancestry informative site to hard calls using a posterior probability threshold of 0.9. Ancestry informative sites that did not have a probability of $\geq 0.9$ for any ancestry state were converted to NA. We summarized per-site and per-tract accuracy for simulated individuals. For sites that were called for the wrong homozygous ancestry state, we counted this as a full error. For sites that incorrectly inferred one haplotype we treated the site as half an error (e.g. the true ancestry state was homozygous parent 1 but the site was called as heterozygous parent 1). These results are summarized in Fig. S7. We provide a script with this release of *ancestryinfer* that will allow users to summarize accuracy for their own simulations following this approach (see Appendix 1; `post_hmm_accuracy_shell_3way.pl`).

Results of our simulations matching the likely hybridization scenario at Tlalica indicated that we expected accuracy to be high in this context (i.e. recent admixture). We note however, that error rates were non-negligible in our simulations of three-way hybrid, especially for heterozygous ancestry tracts. Thus, given these error rates, for analyses presented in the main text, we do not consider individuals inferred to have low-levels of admixture with *X. variatus* (e.g. ~1%) as three-way hybrids.

We also simulated two-way admixture and asked how well the three-way *ancestryinfer* pipeline performs under this scenario. We found that *ancestryinfer* performed well in this scenario, attributing on average 0.6% of genome-wide ancestry the *X. variatus* source population. This result from simulations mirrors our findings in empirical data for *X. birchmanni* x *X. malinche* hybrid populations where *X. variatus* is absent (Fig. S3).

Compared to the previous implementation of *ancestryinfer* that only performed local ancestry inference in hybrids from two source populations, we found that accuracy was more sensitive to choice of priors in the three source population mode (Schumer *et al.*, 2020); Fig. S8). We tested two different scenarios where we misspecified priors: 1) Misspecification of the admixture proportion expected from *X. variatus* and 2) misspecification of the expected time since initial admixture. For the first scenario, we set the prior admixture proportion for *X. variatus* to 0.9 (true value for the simulation = 0.5). For the second scenario we set the prior for the initial admixture between *X. malinche* and *X. birchmanni* to 200 generations before the present and the prior for the initial admixture between this population and *X. variatus* to 20 generations before the present. We found that error rates were substantially higher in both scenarios (1.9% and 2.2% respectively; Fig. S8).

We also found that accuracy was dependent to the specific demographic scenario simulated (Fig. S9). We simulated two scenarios to explore the impact of demographic history on accuracy: 1) admixture between *X. variatus*, *X. birchmanni*, and *X. malinche* as a single pulse, 25 generations before the present (with equal admixture proportions of all three species) and 2) admixture between *X. variatus*, *X. birchmanni*, and *X. malinche* as a single pulse, 50 generations before the present (with equal admixture proportions of all three species). In the first scenario, average error rates increased to 2.3% and in the second scenario average error rates further increased to 3.4% (Fig. S9).

Given the sensitivity of three-way ancestry inference with *ancestryinfer* to misspecification of priors and the exact admixture scenario that we observe, we recommend that users take advantage of the simulation scripts provided with *ancestryinfer* to explore expected accuracy under their scenario of interest. Step-by-step instructions for performing these simulations and setting important parameters are detailed in Appendix 1.

*Supporting Information 2. Additional water quality and chemistry data*

In addition to the metrics reported in the main text, we collected other water quality data, including an alternative approach to quantifying dissolved organic carbon (DOC). We measured concentrations of dissolved organic carbon in water samples that were filtered through a 0.45 μm polyethersulfone membrane (Cat. No. 725-2545, Thermo Fisher Scientific Inc., Waltham, Massachusetts, USA), acidified to pH ~2.0 with HCl, and kept at 4ºC until analysis. Water samples were processed 8-10 days after sampling by the Laboratorio de Ingeniería Ambiental, Instituto de Ingeniería, Universidad Nacional Autónoma de México, using the 720°C catalytic combustion oxidation – non-dispersive infrared detection method with a Total Organic Carbon analyzer (TOC-LCSH, Shimadzu, Kyoto, Kyoto, Japan). We observed the highest levels of DOC at Tlalica, and the lowest levels at Plank (Fig. S5; Table S6). We estimated the concentrations of humic substances based on Malcolm (1991) and Reuter & Perdue (1977). In uncolored freshwater streams, humic substances constitute ~40% of DOC and the fulvic to humic acids ratio is ~9:1, while in organically colored waters humic substances make up ~80% of DOC and the fulvic to humic acids ratio is ~4:1 (Reuter & Perdue, 1977; Malcolm, 1991). Concentrations of humic substances comparable to the estimated concentrations in Tlalica and Calnali Low have been shown to impair chemical and/or visual communication in fish species (Hubbard *et al.*, 2002; Mobley *et al.*, 2020; Table S6). However, the most relevant prior study found no significant effect on *Xiphophorus* mate choice at the estimated humic acid concentrations for Tlalica and Calnali (Fisher *et al.*, 2006; Table S6). This suggests that other water quality parameters may be playing a role in facilitating hybridization among *Xiphophorus variatus* and *X. birchmanni* x *X. malinche*.

We used a 9300 colorimeter (YSI, Yellow Springs, OH) to quantify nitrite. Nitrite has been shown to affect olfaction in other fish species including in *Xiphophorus* (Martinez & Huertas, 2019; Hughes & Huertas, 2022) at concentrations one or two orders of magnitude above nitrite levels measured at Tlalica (Fig. S5). Thus, it is unclear whether nitrite concentrations at Tlalica will affect perception of olfactory cues in *Xiphophorus variatus* and *X. birchmanni* x *X. malinche* hybrids.

We quantified concentrations of dissolved metals in water (filtered using a 0.45 μm polyethersulfone membrane, and acidified to pH ~2.0 with trace metal grade nitric acid) with inductively coupled plasma mass spectrometry (ICP-MS). ICP-MS analysis was carried out by ALS Canada Ltd. following a modified version of the APHA 3030B/EPA 6020A methods (American Public Health Association, 2020; EPA, 1998). Several metals were found at substantially higher concentrations at Tlalica, intermediate levels at Calnali Low and Plaza, and low levels at Plank (Fig. S5). Metals may affect the general integrity and physiology of the olfactory epithelium, target specific types of olfactory sensory neurons, or be taken up by olfactory neurons and affect upper components of the olfactory system (Rouleau *et al.*, 1995; Lazzari *et al.*, 2017; Razmara & Pyle, 2021). Manganese, arsenic, cadmium, aluminum, and zinc were found at substantially higher concentrations at Tlalica (Fig. S5); however, there is little information available about their effects on perception of olfactory cues.

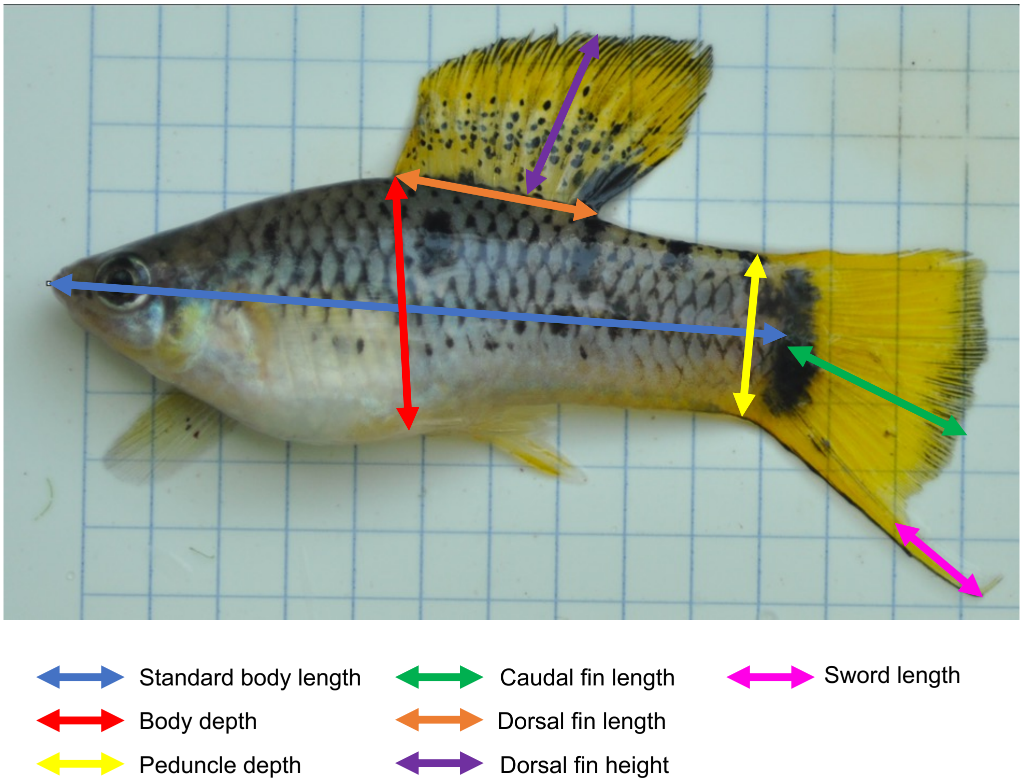


**Fig. S1.** Diagram showing the approach used to measure male phenotypes for PCA of morphological traits in parental species and hybrids.

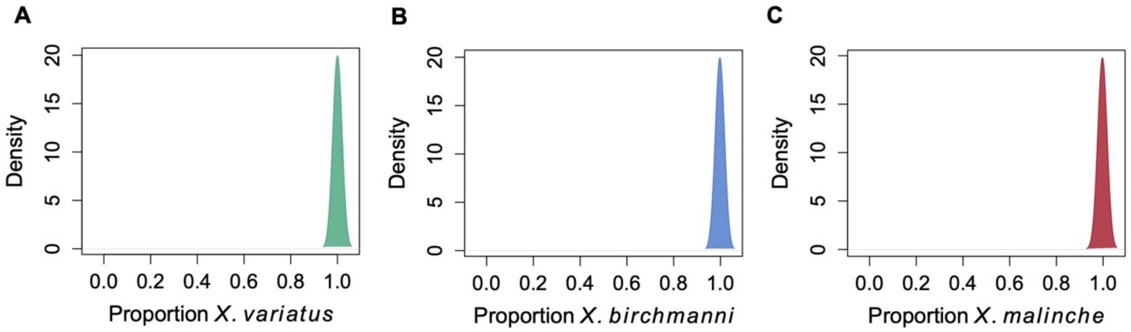


**Fig. S2.** Performance of three-way *ancestryinfer* on pure parental species not used in training or filtering datasets. **A**) Results of inferred genome-wide ancestry for 30 *X. variatus* individuals. **B**) Results for 12 *X. birchmanni* individuals. **C**) Results for 10 *X. malinche* individuals.

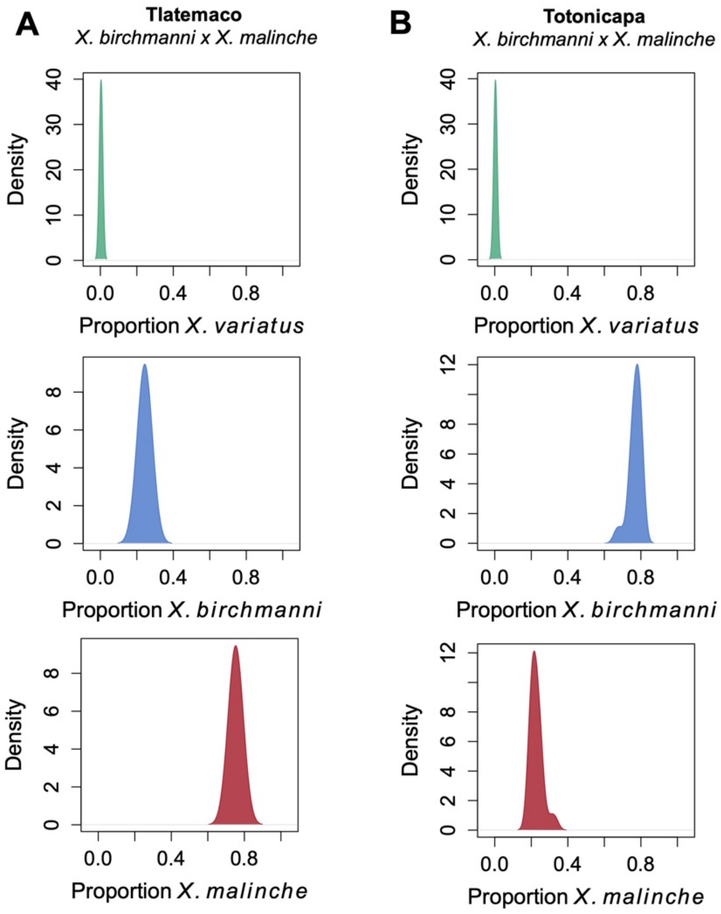


**Fig. S3.** Results of applying *ancestryinfer* in the three source population mode to *X. birchmanni* x *X. malinche* hybrid populations that are not sympatric with *X. variatus*. **A**) Tlatemaco hybrids are inferred to derive <1% of their genome-wide ancestry from *X. variatus* and an average of approximately 75% from *X. malinche* and 24% from *X. birchmanni* respectively. This closely mirrors previous results using 2-way local ancestry inference in this population, see (Schumer *et al.*, 2017). **B**) Totonicapa hybrids are likewise inferred to derive <1% of their genome-wide ancestry from *X. variatus*, 23% from *X. malinche* and 77% from *X. birchmanni*. These results again closely mirror previously reported values for this *X. birchmanni* x *X. malinche* hybrid population (Schumer *et al.*, 2017).

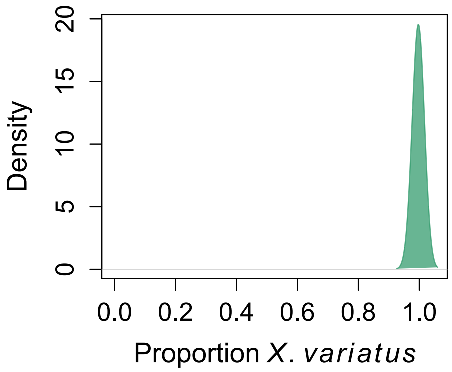


**Fig. S4.** Proportion of the genome inferred to be derived from *X. variatus* in individuals phenotypically determined to be *X. variatus* at the Tlalica site upon collection.

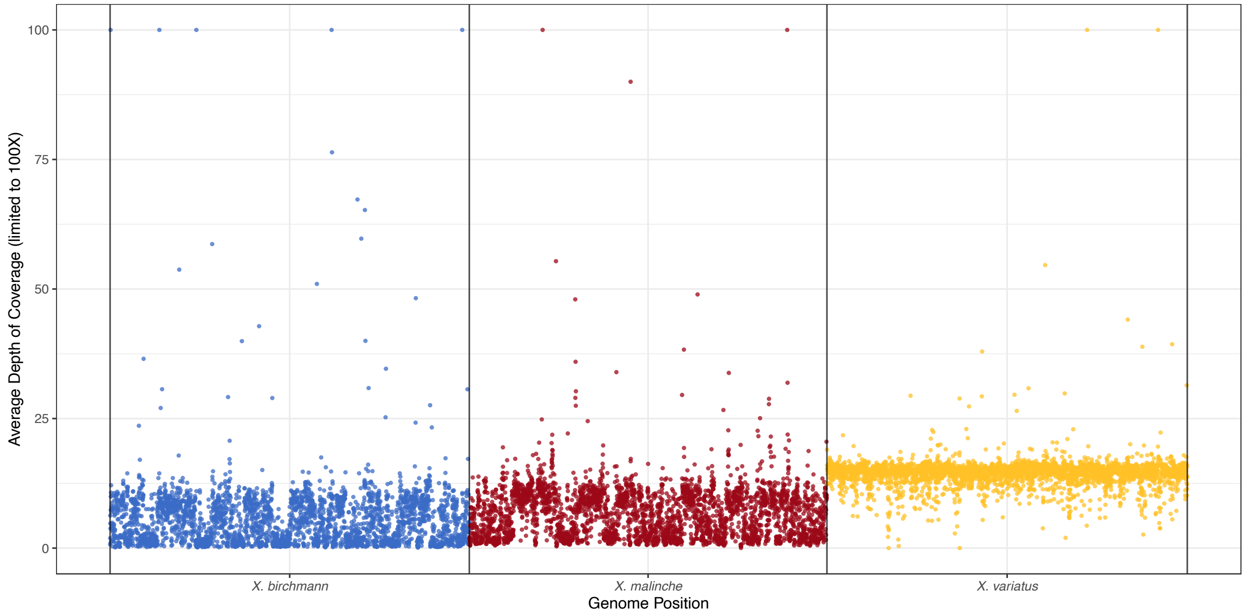


**Fig. S5.** Mean depth of coverage for the *X. birchmanni* (blue), *X. malinche* (red), and *X. variatus* (yellow) genomes for two deep-sequenced individuals that were competitively mapped to reference genomes of all three species using the program sppIDer (Langdon *et al.*, 2018). Coverage averaged in 200 kb windows showed that the *X. variatus* genome was covered at ~15X across almost the whole genome. Coverage across the *X. birchmanni* and *X. malinche* genomes ranged from 0-15X in large tracts, consistent with past recombination between these genomes.

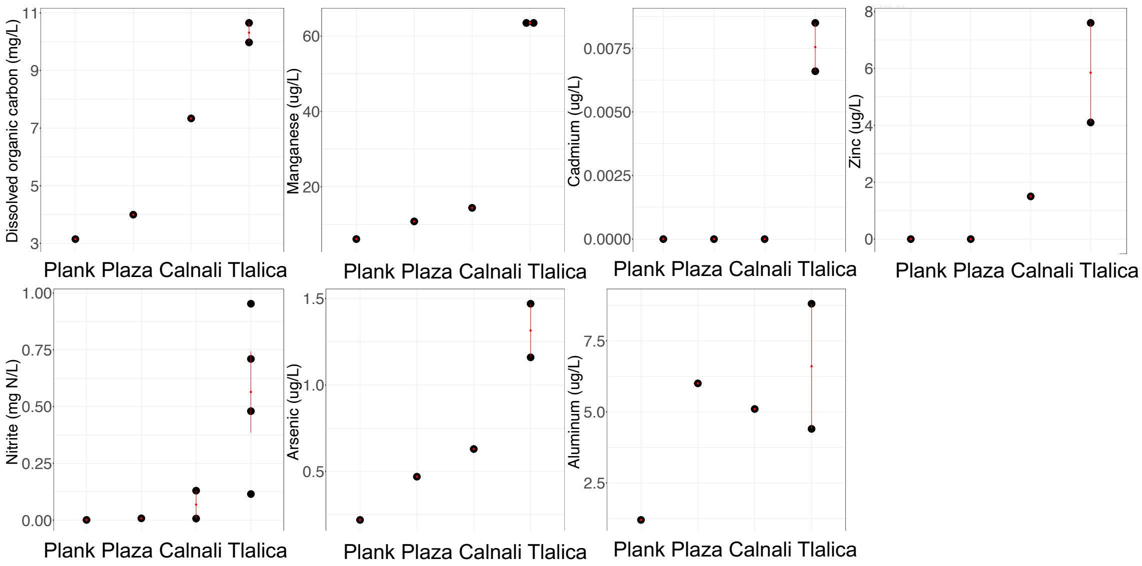


**Fig. S6.** Dissolved organic carbon (mg/L), nitrate (mg N/L), manganese (ug/L), arsenic (ug/L), Cadmium (ug/L), aluminum (ug/L), and zinc (ug/L) levels measured at Plank, Plaza, Calnali Low (Calnali), and Tlalica in May and June of 2022. Black dots represent independent measurements, red dots represent means, and red bars represent one standard error of repeated measurements.

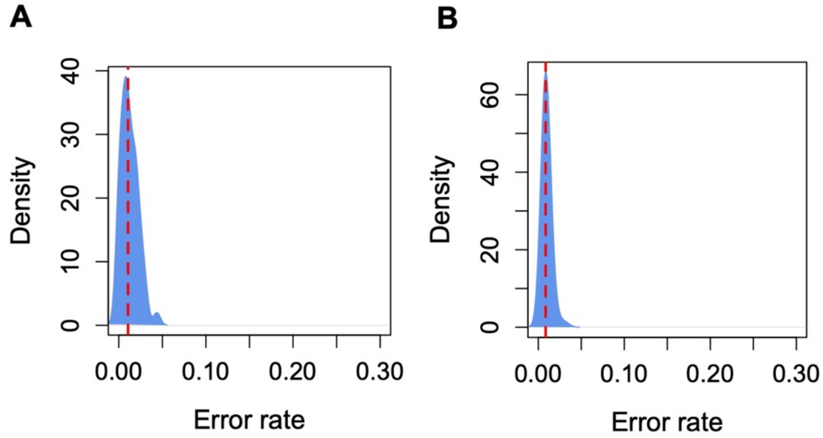


**Fig. S7. A)** Error rates of local ancestry inference in simulations of early generation three-way hybrids between *X. variatus*, *X. birchmanni* and *X. malinche* and **B)** simulations of *X. birchmanni* x *X. malinche* hybrids with parameters matching those observed in the Río Calnali. Plotted here is the error rate in a simulation of 100 diploid individuals with known ancestry. See Supporting Information 1 for more details.

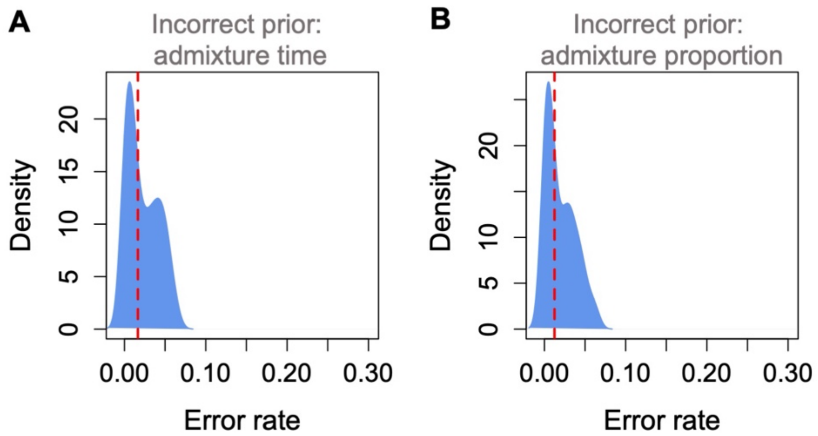


**Fig. S8.** Simulations suggest that the accuracy of three-way local ancestry inference with *ancestryinfer* is sensitive to misspecification of priors for generations since initial admixture and admixture proportions. **A)** Error rates of local ancestry inference in simulations of early generation three-way hybrids formed between *X. variatus*, *X. birchmanni* and *X. malinche* using incorrect priors for time of initial admixture. **B)** Error rates of local ancestry inference in simulations of early generation three-way hybrids formed between *X. variatus*, *X. birchmanni* and *X. malinche* using incorrect priors for initial admixture proportions. Plotted here is the error rate in a simulation of 100 diploid individuals with known ancestry. See Supporting Information 1 for more details.

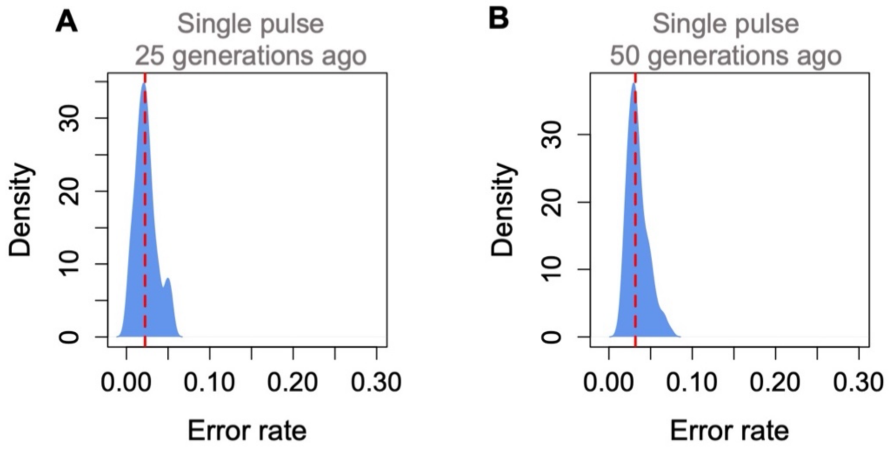


**Fig. S9.** Simulation results suggest that the accuracy of three-way local ancestry inference with *ancestryinfer* is sensitive to the specific demographic scenario simulated. **A)** Error rates of local ancestry inference in simulations of early generation 3-way hybrids formed between *X. variatus*, *X. birchmanni* and *X. malinche* when initial admixture occurred 25 generations between the present (here, equal admixture proportions were simulated). Average error across individuals in this simulation was 2.3% (dashed red line). **B)** Error rates of local ancestry inference in simulations of early generation 3-way hybrids formed between *X. variatus*, *X. birchmanni* and *X. malinche* when initial admixture occurred 50 generations between the present (equal admixture proportions were simulated). Average error across individuals in this simulation was 3.4% (dashed red line). Plotted here is a simulation of 100 diploid individuals with known ancestry. See Supporting Information 1 for more details.

**Table S1**. High coverage sequencing data collected for two individuals that visually appeared to be three-way hybrids.

| **Individual ID** | **Number of 150 bp reads** | **SRA accession** |
|---|---|---|
| CAPS-V-21-M01 | 92,508,957 x 2 | SRAXXXXXXX |
| CAPS-V-21-M02 | 135,901,799 x 2 | SRAXXXXXXX |

**Table S2.** Collection sites, collection years, GPS coordinates, and sample sizes for *X. birchmanni* x *X. malinche* hybrid populations on the Río Calnali. The right column indicates whether *X. variatus* co-occurs with hybrids at each site. Sites are listed in order of decreasing elevation.

| **Collection site (Code – full name)** | **Collection years** | **Coordinates** | **Number of individuals** | ***X. variatus* present at site** |
|---|---|---|---|---|
| CALM – Calnali mid-elevation | 2003, 2004 | 20°53'37.17"N 98°36'36.88"W | 70 | no |
| AGCZ – Aguazarca | 2003, 2016 | 20°53'54.62"N 98°36'7.74"W | 51 | no |
| PILO – Piloncillo | 2017 | 20°53'55.08"N 98°35'17.81"W | 53 | yes |
| PLAZ – Plaza | 2017 | 20°53'49.36"N 98°34'58.86"W | 64 | yes |
| PEAT – Peatonale | 2017 | 20°53'52.97"N 98°34'44.92"W | 55 | yes |
| CALL – Calnali Low | 2004, 2017 | 20°53'57.68"N 98°34'31.58"W | 260 | yes |
| CHAF – Chahuaco falls | 2016, 2017 | 20°54'23.32"N 98°32'14.68"W | 25 | yes |

**Table S3.** Collection years and sites for sequenced individuals from pure *X. birchmanni* and pure *X. variatus* populations.

| **Species** | **Collection site** | **Number of individuals** |
|---|---|---|
| *X. variatus* | COAC | 537 |
| *X. variatus* | CALL | 38 |
| *X. variatus* | Tlalica | 24 |
| *X. birchmanni* | COAC | 122 |
| *X. birchmanni* | GARC | 5 |
| *X. birchmanni* | MAME | 17 |
| *X. birchmanni* | SELV | 2 |

**Table S4.** Principal component loadings of morphological phenotypes of male *X. malinche, X. birchmanni, X. variatus*, *X. birchmanni* x *X. malinche* hybrids and confirmed three-way hybrids.

| | **PC1** | **PC2** | **PC3** | **PC4** | **PC5** | **PC6** | **PC7** |
|---|---|---|---|---|---|---|---|
| **Body length** | 0.808 | 0.163 | 0.333 | 0.444 | -- | -- | -- |
| **Body depth** | 0.355 | -- | -- | -0.681 | 0.577 | -- | -0.264 |
| **Peduncle depth** | 0.22 | -0.454 | -0.192 | -- | 0.139 | -0.286 | 0.776 |
| **Caudal fin length** | 0.21 | -0.789 | -0.117 | -- | -0.242 | 0.332 | -0.388 |
| **Dorsal length** | 0.273 | 0.147 | -- | -0.425 | -0.734 | -0.422 | -- |
| **Dorsal height** | 0.224 | 0.34 | -0.797 | -- | -0.158 | 0.402 | 0.113 |
| **Sword length** | -- | -- | -0.444 | 0.389 | 0.153 | -0.676 | -0.404 |

**Table S5.** Ancestry proportions inferred based on the nuclear genome and mitochondrial genotype for all three-way hybrids identified in this study. M – male, F – female.

| Individual id | Sex | *X. birchmanni* ancestry | *X. malinche* ancestry | *X. variatus* Ancestry | Mitochondrial genotype |
|---|---|---|---|---|---|
| CAPS-V-21-M01 | M | 0.24 | 0.26 | 0.50 | *X. malinche* |
| CAPS-V-21-M02 | M | 0.25 | 0.25 | 0.50 | *X. malinche* |
| swt-CAPS-XI-21-F-1010-L-OGG | F | 0.09 | 0.18 | 0.73 | *X. malinche* |
| swt-CAPS-XI-21-M-1001-UNMRK | M | 0.26 | 0.24 | 0.49 | *X. malinche* |
| CAPS-22-XI-21-F23 | F | 0.26 | 0.25 | 0.49 | *X. malinche* |
| PLAZ-27-F | F | 0.41 | 0.30 | 0.29 | *X. birchmanni* |
| TLLD-V-22-M100 | M | 0.23 | 0.27 | 0.50 | *X. malinche* |
| TLLD-V-22-M101 | M | 0.14 | 0.36 | 0.50 | *X. malinche* |
| CAPS-V-22-F01 | F | 0.14 | 0.36 | 0.50 | *X. malinche* |
| CAPS-V-22-F05 | F | 0.34 | 0.16 | 0.50 | *X. malinche* |
| CAPS-V-22-F07 | F | 0.30 | 0.20 | 0.50 | *X. birchmanni* |
| CAPS-V-22-F08 | F | 0.26 | 0.25 | 0.49 | *X. birchmanni* |
| CAPS-V-22-M9 | M | 0.19 | 0.31 | 0.50 | *X. malinche* |
| CAPS-V-22-M10 | M | 0.25 | 0.25 | 0.50 | *X. malinche* |
| CAPS-V-22-M11 | M | 0.12 | 0.38 | 0.50 | *X. malinche* |
| CAPS-V-22-M12 | M | 0.12 | 0.38 | 0.50 | *X. malinche* |
| CAPS-V-22-M13 | M | 0.20 | 0.30 | 0.50 | *X. malinche* |
| CAPS-V-22-M14 | M | 0.18 | 0.33 | 0.50 | *X. malinche* |

**Table S6.** Estimated concentrations of humic substances based on measured DOC concentrations*.

| | | At 40% of DOC | Fulvic to humic acids (9:1) at 40% of DOC | | | At 80% of DOC | Fulvic to humic acids (4:1) at 80% of DOC | | |
|---|---|---|---|---|---|---|---|---|---|
| Site | DOC (mg C/L) | Humic substances (mg C/L) | Fulvic acid (mg C/L) | Humic acid (mg C/L) | Humic acid (mg/L) | Humic substances (mg C/L) | Fulvic acid (mg C/L) | Humic acid (mg C/L) | Humic acid (mg/L) |
| Plank | 3.15 | 1.26 | 1.13 | 0.13 | 0.25 | 2.52 | 2.02 | 0.50 | 1.01 |
| Calnali Low | 7.34 | 2.93 | 2.64 | 0.29 | 0.59 | 5.87 | 4.70 | 1.17 | 2.35 |
| Tlalica | 10.31 | 4.12 | 3.71 | 0.41 | 0.82 | 8.25 | 6.60 | 1.65 | 3.30 |

* Concentrations of fulvic and humic acids were estimated based on fulvic to humic acids ratios of 9:1 in uncolored freshwater streams and 4:1 in organically colored waters, respectively (Malcolm, 1991; Reuter and Perdue, 1977).
