## Appendix 1 for "Complex hybridization between deeply diverged fish species in a disturbed ecosystem"

### Simulation scripts to model three-way admixture

1) Run SELAM with the desired demographic history for the hybrid population of interest:

```
SELAM -d selam_demography.txt -o selam_simulation_output_parameters_3way.txt -c 2 1 0
```

2) Convert simulations into ancestry tracts and simulated reads with provided scripts:

*Simulation script usage:*

```
perl simulate_3way_hybrids_SELAM.pl genome_species1.fa genome_species2.fa
genome_species3.fa SELAM_results.txt chromosome_length_bp
recombination_rate_per_bp_Morgans chromosome_name number_of_reads output_filename_stem
```

*Example:*

```
perl simulate_3way_hybrids_SELAM.pl Xbirchmanni_COAC_ pseudoref.fasta
Xmalinche_CHIC_pseudoref.fasta Xvariatus_COAC_pseudoref.fasta
admix_simulation_demography_output_ 3way.txt 25000000 0.00000002 ScyDAA6-1508-HRSCAF-1794
100000 admix_SELAM_sim_3way
```

Note that the reference genomes need to be in the same coordinate space to use this simulation script. The simulated reads will be derived from the focal chromosome only.

*Output*:

This script will generate paired-end reads for each individual, a fasta file for each individual, and the true ancestry along the chromosome for each haplotype. For example, for the first female simulated by SELAM the files produced by the above command will be:

```
admix_SELAM_sim_3way_1_sex1.fa

admix_SELAM_sim_3way_1_sex1_r1.fq.gz
admix_SELAM_sim_3way_1_sex1_r2.fq.gz

admix_SELAM_sim_3way_1_sex1_tracts_hap1.bed
admix_SELAM_sim_3way_1_sex1_tracts_hap2.bed
```

For downstream analysis (step 3), you will need to make a text file with the true ancestry states for each haplotype for each individual. The format of this file should be:

```
admix_SELAM_sim_3way_1_sex1_tracts_hap1.bed\tadmix_SELAM_sim_3way_1_sex1_tracts_hap2.bed\n
admix_SELAM_sim_3way_2_sex1_tracts_hap1.bed\tadmix_SELAM_sim_3way_2_sex1_tracts_hap2.bed\n
```

3) Run *ancestryinfer* (see below) and summarize accuracy results:

*Accuracy script usage:*

```
perl post_hmm_accuracy_shell_3way.pl ancestry_results_file_stem
list_of_true_ancestry_beds posterior_probability_threshold directory
```

*Example:*

```
perl post_hmm_accuracy_shell_3way.pl allchrs.tsv admixture_SELAM_sim_bed_list 0.9 ./
```

### ***ancestryinfer*** **three-way pipeline**

Data can be input into the *ancestryinfer* pipeline to run local ancestry inference following (Corbett-Detig & Nielsen, 2017).

#### Install

*Option 1 – install dependencies:*

```
git clone https://github.com/Schumerlab/ancestryinfer.git
```

To install dependencies, follow instructions outlined in:

```
installation_instructions.txt
```

Test that the install and pipeline are working:

```
cd ancestryinfer
```

```
perl Ancestry_HMM_parallel_v7.pl hmm_configuration_file_nonparallel.cfg
```

*Option 2 – load docker file for dependencies:*

```
docker pull schumer/mixnmatch-ancestryinfer-image:mixnmatch-ancestryinfer-docker
```

```
docker run -it mixnmatch-ancestryinfer-image bash
```

#### Setting parameters in the configuration file

There is are example configuration files available on github.

Example files for 2-way local ancestry inference:

```
hmm_configuration_file_parallel.cfg
```

`hmm_configuration_file_nonparallel.cfg`

Example files for 3-way local ancestry inference:

`hmm_configuration_file_3way_parallel.cfg`

`hmm_configuration_file_3way_nonparallel.cfg`

Parameter descriptions:

| Parameter | Description | Example | Include if |
|---|---|---|---|
| genome1= | User provided fasta file for species 1 | genome1=xiphophorus_birchmanni_10x_12Sep2018_yDAA6.fasta | Always |
| genome2= | User provided fasta file for species 2 | genome2=Xmalinche_dovetail_assembly.fa | Always |
| genome3= | User provided fasta file for species 3 | Genome3=Xvariatus_10x_assembly.fa | Required if performing 3 way ancestry calling |
| read_type= | Indicated whether data is paired end or single end | read_type=PE | Always |
| read_list= | Provide list (including full paths) to the reads to be analyzed | read_list=combined_all_call_hybrids_read_list<br><br>example list format for paired end data (single end file should contain one line per individual):<br><br>./reads/CALL1_read1.fq.gz\t./reads/CALL1_read2.fq.gz<br>./reads/CALL2_read1.fq.gz\t./reads/CALL2_read2.fq.gz<br>./reads/CALL3_read1.fq.gz\t./reads/CALL3_read2.fq.gz | Always |
| read_length= | Provide expected read length. If read lengths of input samples differ, use the longest read lengths. | read_length=150 | Always |
| mapping_quality= | Required mapping quality for a read to be retained | mapping_quality=20 | Optional, if not specified the mapping quality threshold used is 30 |
| prop_genome_genome1_parent= | Expected proportion of the genome derived from the parent | prop_genome_genome1_parent=0.5 | If not provided, AncestryHMM will attempt to estimate (may |

| | | | |
|---|---|---|---|
| | species listed under genome1 | | increase run time) |
| prop_genome_genome2_parent= | Expected proportion of the genome derived from the parent species listed under genome2 | prop_genome_genome2_parent=0.5 | If not provided, AncestryHMM will attempt to estimate if 3 way ancestry calling is being performed (may increase run time) |
| number_indiv_per_job= | Parallelize jobs such that each job processes this number of individuals.<br><br>Low numbers mean high parallelization and high number mean low parallelization. | number_indiv_per_job=1 | Always |
| program_path= | Path to the program install folder | program_path=/home/groups/schumer/shared_bin/Ancestry_HMM_pipeline | If not provided the program will assume necessary scripts and programs are in the working directory |
| provide_AIMs= | Coordinates and identities of ancestry informative sites that distinguish the two parent species | provide_AIMs=Xbirchmanni10xgenome_ancestry_informative_sites_filterF1<br><br>Example list format for 2 or 3-way ancestry calling:<br><br>ScyDAA6-2-HRSCAF-26 58345 T C<br>ScyDAA6-2-HRSCAF-26 58976 T A<br>ScyDAA6-2-HRSCAF-26 59896 T C<br>ScyDAA6-2-HRSCAF-26 60164 G A<br>ScyDAA6-2-HRSCAF-26 63105 G A<br>ScyDAA6-2-HRSCAF-26 65532 G A<br>ScyDAA6-2-HRSCAF-26 66290 C A<br>ScyDAA6-2-HRSCAF-26 68233 T C<br>ScyDAA6-2-HRSCAF-26 70398 G A<br>ScyDAA6-2-HRSCAF-26 73869 G A | Required unless provided genomes are on the same coordinate system and can be auto detected<br><br>*Always required for 3 way ancestry calling* |
| provide_counts= | Counts of parental allele frequencies at ancestry informative sites (and recombination rates between | provide_counts=Xbirchmanni10xgenome_Xmalinche_observed_parental_counts_filterF1<br><br>Example format for 2-way ancestry calling:<br><br>ScyDAA6-2-HRSCAF-26 163722 129 3 0 54 0.00000078<br>ScyDAA6-2-HRSCAF-26 166158 135 5 0 54 0.00001374 | If not provided, the program will assume that provided ancestry informative sites are fixed between species |

| | | | |
|---|---|---|---|
| | adjacent sites if available) | ScyDAA6-2-HRSCAF-26 166535 6 0 0 6 0.00000754<br><br>Columns are:<br><br>Chromosome site allele1_count_parent1 allele2_count_parent1 allele1_count_parent2 allele2_count_parent2 recombination_rate<br><br>Example format for 3-way ancestry calling:<br><br>ScyDAA6-2-HRSCAF-26 230490 12 0 0 11 94 0 4.1e-06<br>ScyDAA6-2-HRSCAF-26 240248 220 6 0 54 74 0 8.78e-06<br>ScyDAA6-2-HRSCAF-26 240783 218 6 0 54 59 0 4.26e-06<br>ScyDAA6-2-HRSCAF-26 241074 217 5 0 54 27 0 5.82e-06<br><br>Chromosome site allele1_count_parent1 allele2_count_parent1 allele1_count_parent2 allele2_count_parent2 allele1_count_parent3 allele2_count_parent3 recombination_rate | *Always required for 3 way ancestry calling* |
| per_site_error = | Per-site error parameter for HMM (i.e. due to sequencing error, contamination, etc) | per_site_error=0.02 | Always |
| gen_initial_admix_p1_p2= | Estimated generation of initial admixture of genome 1 and genome 2 species | gen_initial_admix_p1_p2=20 | If not provided, AncestryHMM will attempt to estimate (may increase run time) |
| gen_initial_admix_p3= | Estimated generation of initial admixture of genome 3 species | gen_initial_admix_p3=60 | If not provided and 3 way admixture is indicated, AncestryHMM will attempt to estimate (may increase run time) |
| focal_chrom_list= | Provide a list of chromosomes to run (other chromosomes will not be run) | focal_chrom_list=mychrs.txt<br><br>Example:<br><br>ScyDAA6-2-HRSCAF-26<br>ScyDAA6-7-HRSCAF-50 | Not required |
| rec_M_per_bp= | Estimated recombination | rec_M_per_bp=0.00000002 | Always; Use an estimate for a |

| | | | |
|---|---|---|---|
| | rate in Morgans/bp | | related species if not available |
| max_alignments = | Limit analysis to a maximum number of alignments (for computational speed) | max_alignments=2000000 | Optional |
| retain_intermediate_files= | Keep all intermediate files. Warning: setting this to 1 results in a high space footprint for a large run; only recommended for troubleshooting. | retain_intermediate_files=0 | Options are 1 to keep or 0 to delete. |
| posterior_thresh= | Posterior probability threshold to use for identifying ancestry transition intervals | posterior_thresh=0.9 | Recommended 0.8-1 |
| job_submit_command= | Option to run sequentially if using Docker image for dependencies or from a desktop computer. Set bash to run sequentially and sbatch to run in parallel on a slurm cluster | job_submit_command=bash<br><br>or<br><br>job_submit_command=sbatch | Always required |
| slurm_command_map=<br><br>slurm_command_variant_call=<br><br>slurm_command_hmm= | If running on a slurm cluster, provide cluster specific parameters for queues, time & memory | slurm_command_map=#!/bin/sh #SBATCH --ntasks=1 #SBATCH --cpus-per-task=1 #SBATCH -p schumer --mem=64000 #SBATCH --time=02:30:00<br><br>slurm_command_variant_call=#!/bin/sh #SBATCH --ntasks=1 #SBATCH --cpus-per-task=1 #SBATCH -p schumer --mem=64000 #SBATCH --time=05:00:00<br><br>slurm_command_hmm=#!/bin/sh #SBATCH --ntasks=1 #SBATCH --cpus-per-task=1 #SBATCH -p schumer --mem=64000 #SBATCH --time=03:00:00 | Required if running on a cluster |

### Examples

Several example files are available with the git repository including example configuration files

**Running the pipeline**

After setting the parameters in the configuration file and loading required dependencies, simply run:

```
perl $PATH/Ancestry_HMM_parallel_v7.pl hmm_configuration_file_3way_parallel.cfg
```

where $PATH is the path to your simulator install (e.g. `/home/groups/schumer/shared_bin/Ancestry_HMM_pipeline`)
